# Brain cell type specific proteomics approach to discover pathological mechanisms in the childhood CNS disorder mucolipidosis type IV

**DOI:** 10.1101/2023.05.04.539472

**Authors:** Madison Sangster, Sanjid Shahriar, Zachary Niziolek, Maria Carla Carisi, Michael Lewandowski, Bogdan Budnik, Yulia Grishchuk

## Abstract

1 Mucolipidosis IV (MLIV) is an ultra-rare, recessively inherited lysosomal disorder resulting from inactivating mutations in *MCOLN1*, the gene encoding the lysosomal cation channel TRPML1. The disease primarily affects the central nervous system (CNS) and manifests in the first year with cognitive and motor developmental delay, followed by a gradual decline in neurological function across the second decade of life, blindness, and premature death in third or fourth decades. Brain pathology manifestations in MLIV are consistent with hypomyelinating leukodystrophy with brain iron accumulation. Presently, there are no approved or investigational therapies for MLIV, and pathogenic mechanisms remain largely unknown. The MLIV mouse model, *Mcoln1*^*-/-*^ mice, recapitulates all major manifestations of the human disease. Here, to better understand the pathological mechanisms in the MLIV brain, we performed cell type specific LC-MS/MS proteomics analysis in the MLIV mouse model and reconstituted molecular signatures of the disease in either freshly isolated populations of neurons, astrocytes, oligodendrocytes, and neural stem cells, or whole tissue cortical homogenates from young adult symptomatic *Mcoln1*^*-/-*^ mice. Our analysis confirmed on the molecular level major histopathological hallmarks of MLIV universally present in *Mcoln1*^*-/-*^ tissue and brain cells, such as hypomyelination, lysosomal dysregulation, and impaired metabolism of lipids and polysaccharides. Importantly, pathway analysis in brain cells revealed mitochondria-related alterations in all *Mcoln1*^*-/-*^ brain cells, except oligodendrocytes, that was not possible to resolve in whole tissue. We also report unique proteome signatures and dysregulated pathways for each brain cell population used in this study. These data shed new light on cell-intrinsic mechanisms of MLIV and provide new insights for biomarker discovery and validation to advance translational studies for this disease.

## 2 Introduction

Mucolipidosis type 4 (MLIV) is a lysosomal disease that affects the central nervous system and is inherited in an autosomal-recessive manner. The typical form of MLIV, caused by the absence of functional TRPML1 protein or its complete loss of function, is characterized by hypomyelinating leukodystrophy with brain iron accumulation and manifests with severely impaired psychomotor development and a gradual neurological decline paralleled by cerebellar neurodegeneration and neuroaxonal injury (1, 2, 3). Retinal dystrophy also develops in the first years of life and leads to blindness by the second decade. Brain MRI abnormalities in childhood include marked hypoplasia of the corpus callosum, decreased subcortical white matter, ferritin deposition in the basal ganglia, and relative preservation of cortical gray matter. Later in the course of disease, over the first two decades, the cerebellum degenerates, and, eventually, diffuse cerebral atrophy may become apparent (2, 3).

Although it is not fully understood how early brain development is affected in MLIV, subcortical hypomyelination and iron accumulation in the basal ganglia have been reported in MLIV at gestational stage (4). Despite this early evidence of brain pathology, patients generally achieve their developmental motor milestones up to six months of age and present with axial hypotonia and delayed gross motor development around one year of age (5, 6). Corneal opacities often present from birth, elevated plasma gastrin, and delayed neuromotor development in infants warrant genetic testing and are suggestive of MLIV. Most patients do not achieve independent ambulation; however, some are able to crawl in an uncoordinated fashion and take steps with a walker that provides full truncal support. Impaired finger articulation limits fine motor function, and most patients will only develop an inferior or modified pincer grasp (5, 7). Motor development in MLIV is negatively impacted by both pyramidal and extrapyramidal tract dysfunction. The pyramidal signs, such as spasticity, diffuse hyperreflexia, and decreased force with volitional muscle activation gradually increase in severity across the lifetime. The extrapyramidal motor dysfunction manifests as rigidity, dystonic posturing, postural instability, bradykinesia, tremor, truncal ataxia, and jerk nystagmus. These signs are typically present in the first year of life, but progressively worsening spasticity makes the presence of extrapyramidal signs difficult to discern later in life, and their progression remains unstudied.

Cognitive development in patients with MLIV is difficult to assess due to motor, expressive language, and visual impairment. Language development is impaired due to a combination of oral motor dysfunction and a poorly defined cognitive component. Patients can follow simple and multi-step commands, laugh at jokes, and exhibit appropriate social responses to verbal context. Verbal communication is typically limited to fewer than 10 understandable words, but many patients use hand signs and adaptive communication devices.

Due to the extremely limited availability of MLIV human brain tissue for research, the MLIV mouse model has become an important resource for studying underlying mechanism of diseases and investigating the role of TRPML1 in brain development and myelination. Mice with *Mcoln1* knock out exhibit all major symptoms of MLIV disease, including cognitive and motor dysfunction, brain and eye pathology, and elevated plasma gastrin (8, 9, 10, 11). *Mcoln1*^*-/-*^ mice appear indistinguishable from their wild-type and *Mcon1*^*-/+*^ littermates at birth but develop cognitive and motor deficits by two months, with the progression of motor dysfunction evident through rotarod and balance beam tests. Motor deficits progress to rear limb paralysis by seven months, and premature death occurs a few weeks later (11), (12).

Brain histopathology in *Mcoln1*^*-/-*^ mice shows typical characteristics seen in other mouse models of lysosomal disorders, such as an accumulation of autofluorescent material, neuronal accumulation of gangliosides and cholesterol, and increased autophagy substrate P62/SQSTRM, indicating inhibited autophagic and lysosomal function. All human brain pathology hallmarks of MLIV such as decreased myelination, glial activation and partial Purkinje neurons loss in late-stage cerebellum are also present in *Mcoln1*^*-/-*^ mice (12, 13). Brain tissue analysis of younger *Mcoln1*^*-/-*^ mice shown that many of the pathological hallmarks present in the brain at the terminal stage, such as activation of glia, reduced myelination, and lysosomal material accumulation, are already present as early as at post-natal day 10, before onset of the first detectable motor dysfunction in this model that develops by 2 months of age [12]. Remarkably, *Mcoln1*^*-/-*^ mice do not show significant brain atrophy or neuronal loss throughout the course of the disease, except for the partial loss of Purkinje cells in the cerebellum mentioned earlier (9, 12). These findings, along with the data from MLIV patients, suggest that TRPML1 plays an essential role in early brain development, and that loss of its function leads to widespread brain pathology and dysfunction of various brain cell types but does not cause overt neurodegeneration.

Overall, histological analysis of *Mcoln1*^*-/-*^ brain and studies on isolated cells indicate that all brain cell types are affected by loss of TRPML1 function (8, 9), yet resolution of these changes on the molecular level and understanding of the functional impact of TRPML1 loss in major brain cell types is missing. Here, we set out to reconstitute the proteomic profiles of whole cortical tissue and freshly isolated neurons, neural stem cells, oligodendrocytes, microglia, and astrocytes from *Mcoln1*^*-/-*^ and control littermates at the early symptomatic stage of disease at 3 months.

## Materials and Methods

### Animals

*Mcoln1*^-/-^ mice (RRID:IMSR_JAX:027110) were maintained as previously described (11). Genotyping was performed by Transnetyx using real-time qPCR (www.transnetyx.com). The *Mcoln1*^+/-^ breeders for this study were obtained by backcrossing onto a C57BL/6J background for more than 10 generations. Experimental cohorts were obtained from *Mcoln1*^+/-^ x *Mcoln1*^-/-^ mating. *Mcoln1*^+/-^ and *Mcoln1*^+/+^ littermates were used as controls. Experiments were performed according to the Institutional and National Institutes of Health guidelines and approved by the Massachusetts General Hospital Institutional Animal Care and Use Committee. Female mice were used at 3 months of age.

### Brain tissue dissociation

Brain tissue dissociation was performed according to 31551601 with some modifications. Immediately after euthanasia using a carbon dioxide chamber, the brain was extracted, the meninges removed, and the dissected cortex was placed in ice-cold HBSS buffer (no calcium, no magnesium, 1% GlutaMAX, 5% trehalose). After a short spin-down and removal of supernatant, the Adult Brain Dissociation kit (Miltenyi #130-107-677, RRID:SCR_020295) was used with a few modifications to obtain a single cell solution. In enzyme mix 1, the papain concentration was halved and 5% trehalose (Sigma-Aldrich, #T0167) was added; 45uM actinomycin D (Sigma-Aldrich #SBR00013) was added to both enzyme mixes. The samples were incubated under continuous rotation at 34^0^C. Each enzyme mix was added separately in between incubations. The cortices were dissociated manually using a serological pipet followed by a fire-polished Pasteur pipette. 10% ovomucoid protease inhibitor (Worthington Biochemical #LK003182) was added to quench the reaction. The solution was washed with a wash buffer comprised of 0.5% BSA (Miltenyi #130-091-376), 1% GlutaMAX (Gibco # 35050061) in HBSS with calcium and magnesium (Gibco # 14185052), and cell clumps were removed with a 70um cell strainer (Miltenyi #130-098-462), then washed two more times to remove myelin and debris. Erythrocytes were removed with the Miltenyi red blood cell removal solution diluted in ddH_2_O and a 5-minute incubation at 4°C, followed by an additional wash and spin down.

### Cell labelling and flow cytometry

The cells were resuspended in ice-cold labelling buffer (0.1% BSA, 2mM EDTA, 1% GlutaMAX, and 5% trehalose in HBSS without calcium or magnesium) and incubated with FcR blocking reagent (Miltenyi #130-092-575) for 10 minutes at 4°C in the dark under continuous rotation to reduce non-specific cell labelling. The cells were then labelled with the following antibodies: anti-ASCA-2-APC (Miltenyi #130-117-535, RRID:AB_2727978; 1:14) for astrocytes, anti-O4-488 (R&D #FAB1326G, RRID:AB_2936942; 1:27) for oligodendrocytes, anti-CD200-PE (Biolegend #123808, RRID:AB_2073942;1:90) for neurons, anti-CD11b-BV510 (BD #562950, RRID:AB_2737913; 1:90) for microglia, and anti-CD15-PerCP-eFluor710 (eBioscience #46-8813-42, RRID:AB_11217476; 1:22.5) for neuronal stem cells for 15 minutes at 4°C in the dark under continuous rotation. The cells were washed, pelleted, and resuspended in 15ml ice-cold FACS buffer (0.5% BSA, 1% GlutaMAX, and 5% trehalose in HBSS with calcium and magnesium) per cortex. Cells were incubated with 10μM Dyecycle Violet (Thermo V35003) in the FACS tubes 15 min prior sorting to sort out cellular debris and dead cells. Live nucleated Dyecycle Violet + cells were then sorted with a Moflo Astrios instrument (Beckman Coulter, RRID:SCR_018893) with a 100 μm nozzle at 24 psi. Gates were set manually with compensation beads (Life Technologies, A10497) and appropriate control samples, and data were analyzed with FlowJo software (v.10).

### Individual Brain Cell Specific Sample Preparation

FACS sorted astrocytes, oligodendrocytes, neurons, microglia, and NSC were stored in -80°C in separate 2mL Eppendorf tubes. Frozen samples were placed on Corning LSE Digital Dry Bath at 95°C for 5min. Samples were immediately placed on ice bath for 5 min. **Digestion:** A stock solution of Trypsin Platinum, Mass Spec Grade (Promega) was made up at 100ug/mL in 50mM TEAB. A specific volume of stock trypsin was added to each sample vial for each set of brain cells using a 1:50 ratio (Trypsin:Protein). Samples were incubated for 2 hours at 50°C and shaken/mixed at 350rpm on Eppendorf ThermoMixer C. **TMT Labeling:** Samples, now peptides, were labeled using 4uL of TMTpro Mass Tag Labels (ThermoScientific). After labeling, samples were then placed on Eppendorf ThermoMixer C for 45min and shaken/mixed for 45min; this ensures labels are covalently bonded to peptides. Labeling reaction was quenched for 10min using 1uL of 5% hydroxyalamine. After quenching, samples were pooled into one 2mL Eppendorf tube and then dried down using Eppendorf Vacufuge plus. **Desalting Samples:** Dried samples were resuspended in 300uL of 0.1% TFA in ultrapure HPLC grade water and vortexed to ensure full solubility. Samples were then desalted using Pierce™ Peptide Desalting Spin Columns (ThermoScientific). Final eluate (desalted samples) contained 600uL of 50% HPLC grade Acetonitrile & 50% ultrapure HPLC grade water. Desalted samples were then transferred to HPLC vials and were dried down using Eppendorf Vacufuge plus. Dried and desalted samples were resuspended in 6uL of 0.1% Formic Acid in ultrapure HPLC grade water and were then injected on LC-MS/MS.

### Brain tissue preparation

Brain tissue was stored in -80°C in 2mL Eppendorf tubes **Protein Extraction Protocol:** Samples were placed on dry ice and frozen brain tissue was ground up inside tube using pestle. 130uL of 5% SDS was then added to ground up tissue. Samples were centrifuged at 15,000g for 5min on Eppendorf Centrifuge 5420. Supernatant (100uL) was then transferred to new 2mL Eppendorf tubes. The remaining 30uL was stored in -80°C for future work. **Reduction/Alkylation and Acidification:** 8.7uL of 10mM TCEP was added to each sample. Samples were then shaken/mixed for 30 minutes at 350 rpm on Eppendorf ThermoMixer C. 8.7uL of 10mM IAA was added to reduced samples.

Samples were then shaken/mixed for 10 minutes at 350 rpm on Eppendorf ThermoMixer C. 21.7uL of 12.5% Phosphoric Acid was added to samples. **Trap and Clean Protein Protocol:** 1522uL of Binding/Wash Buffer (100mM TEAB in 90% Methanol) was added to sample and immediately vortexed; sample visually appeared as milky colloidal suspension. Colloidal suspension was trapped by transferred to S-Trap mini column and centrifuged at 4000g for 30seconds on Eppendorf Centrifuge 5420; flow through discarded. Protein was cleaned on S-Trap mini column (Protifi) by washing with 1522uL Binding/Wash Buffer and centrifuged at 4000g for 30seconds on Eppendorf Centrifuge 5420; this was repeated for total of 3X and each flow through was discarded. 3X cleaning in previous step is critical as it ensures SDS is washed from protein sample. **Digestion:** A stock solution of Trypsin Platinum, Mass Spec Grade (Promega) was made up at 100ug/mL in 50mM TEAB. A specific volume of stock trypsin was added directly to the top of each S-Trap mini column using a 1:50 ratio (Trypsin:Protein). Samples were incubated for 2 hours at 50°C and shaken/mixed at 350rpm on Eppendorf ThermoMixer C. **Elute Peptides:** Elution 1: After digest was completed 80uL of 50mM TEAB in water (pH 8.5) was added directly to each S-Trap mini column and then centrifuged at 4000g for 1 minute on Eppendorf Centrifuge 5420; flow through containing peptides was set aside. Elution 2: 80uL of 0.2% Formic Acid added directly to each S-Trap mini column and then centrifuged at 4000g for 1 minute on Eppendorf Centrifuge 5420; flow through containing peptides was set aside. Elution 3: 80uL of 50% Acetonitrile in water was added directly to each S-Trap mini column and then centrifuged at 4000g for 1 minute on Eppendorf Centrifuge 5420; flow through containing peptides was set aside. For each sample, the 3 elution’s were combined and set aside for TMT labeling. **TMT Labeling:** Samples, now peptides, were labeled using 40uL of TMTpro Mass Tag Labels (ThermoScientific). After labeling, samples were then placed on Eppendorf ThermoMixer C for 45min and shaken/mixed for 45min; this ensures labels are covalently bonded to peptides. Labeling reaction was quenched for 10min using 8uL of 5% hydroxyalamine.

After quenching, samples were pooled into one 2mL Eppendorf tube and then dried down using Eppendorf Vacufuge plus. **Fractionation:** Dried samples were resuspended in 120uL of 0.1% TFA in ultrapure HPLC grade water and vortexed to ensure full solubility. Samples were then fractionated using Pierce™ High pH Reversed-Phase Peptide Fractionation Kit Columns (ThermoScientific).

Each fraction contained 120uL of volume for a total of 20 fractions. Each fraction was centrifuged at 3000g for 2minon Eppendorf Centrifuge 5420; flow throughs collected and transferred to HPLC vials. All fractions were dried down using Eppendorf Vacufuge plus and resuspended in 6uL of 0.1% Formic Acid in ultrapure HPLC grade water and were then injected on LC-MS/MS.

### Mass spectrometry analysis

After separation each fraction was submitted for single LC-MS/MS experiment that was performed on an Exploris 240 Orbitrap (Thermo Scientific, RRID:SCR_022216) equipped with NEO (Thermo Scientific) nanoHPLC pump. Peptides were separated onto a 300 μm × 5 mm PepMap C18 trapping column (Thermo Scientific, Lithuania) followed by DNV PepMap Neo 75umx150mm analytical column (Thermo Scientific, Lithuania). Separation was achieved through applying a gradient from 5–25% ACN in 0.1% formic acid over 120 min at 250 nl min−1. Electrospray ionization was enabled through applying a voltage of 1.8 kV using a PepSep electrode junction at the end of the analytical column and sprayed from stainless still PepSep emitter SS 30μm LJ (Odense, Denmark). The Exploris Orbitrap was operated in data-dependent mode for the mass spectrometry methods. The mass spectrometry survey scan was performed in the Orbitrap in the range of 450 –900 m/z at a resolution of 1.2 × 10^5^, followed by the selection of the ten most intense ions (TOP10) ions were subjected to HCD MS2 event in Orbitrap part of the instrument. The fragment ion isolation width was set to 0.8 m/z, AGC was set to 50,000, the maximum ion time was 150 ms, normalized collision energy was set to 34V and an activation time of 1 ms for each HCD MS2 scan.

### Mass spectrometry data analysis

Raw data were submitted for analysis in Proteome Discoverer 3.0.1.23 (Thermo Scientific, RRID:SCR_014477) software with Chimerys. Assignment of MS/MS spectra was performed using the Sequest HT algorithm and Chimerys (MSAID, Germany) by searching the data against a protein sequence database including all entries from the Mouse Uniprot database (SwissProt 19,768 2019; RRID:SCR_002380) and other known contaminants such as human keratins and common lab contaminants. Sequest HT searches were performed using a 20 ppm precursor ion tolerance and requiring each peptides N-/C termini to adhere with Trypsin protease specificity, while allowing up to two missed cleavages. 18-plex TMT tags on peptide N termini and lysine residues (+304.207146 Da) was set as static modifications and Carbamidomethyl on cysteine amino acids (+57.021464 Da) while methionine oxidation (+15.99492 Da) was set as variable modification. A MS2 spectra assignment false discovery rate (FDR) of 1% on protein level was achieved by applying the target-decoy database search. Filtering was performed using a Percolator (64bit version, reference 1). For quantification, a 0.02 m/z window centered on the theoretical m/z value of each of the six reporter ions and the intensity of the signal closest to the theoretical m/z value was recorded. Reporter ion intensities were exported in result file of Proteome Discoverer 3.0 search engine as an excel tables. The total signal intensity across all peptides quantified was summed for each TMT channel, and all intensity values were adjusted to account for potentially uneven TMT labeling and/or sample handling variance for each labeled channel.

Proteomics spectral matching (PSM) level data obtained from each TMT channel was subjected to comprehensive analysis in R. The datasets were preprocessed by first imputing missing values with random forest, followed by removal of PSMs that had no data on isolation interference, or showed isolation interference values greater than 70, or were mapped to multiple proteins or no proteins at all. The preprocessed PSM level data was then aggregated to the protein level and normalized using the variance stabilizing normalization method. Differential expression analysis was performed using the R limma package to compare the *Mcoln1*^*-/-*^ mouse brain tissue/cell types against their control counterparts. To visualize the results, volcano plots were generated for each dataset, emphasizing proteins that were up- or downregulated 1.5-fold in the mutants relative to the wild-type samples, and were significantly different at p values < 0.1. The overlap in significantly up- and downregulated proteomic signatures among the five brain cell types and brain tissue were visualized using Venn Diagrams (https://bioinformatics.psb.ugent.be/webtools/Venn/). Lastly, the Database for Annotation, Visualization and Integrated Discovery (DAVID; RRID:SCR_001881) Bioinformatics functional annotation tool was used to identify signaling pathways, metabolic pathways and protein interactions enriched in *Mcoln1*^*-/-*^ brain tissue and cell types relative to their control counterparts. Pathways with FDR ≤ 0.05 were deemed to be significantly altered.

## 3 RESULTS

To resolve cell type specific proteomes in adult mouse brain in symptomatic *Mcoln1*^*-/-*^ (n=5) and control mice (n=5), pieces of cortical tissue were individually homogenized and cells were freshly isolated using an optimized protocol detailed in Material and Methods, labeled with cell type specific fluorophore-conjugated antibodies (anti-ASCA-2 for astrocytes, anti-O4 for oligodendrocytes, anti-CD200 for neurons, anti-CD11b for microglia, and anti-CD15 for neuronal stem cells), collected using fluorescence activated cell sorting (FACS), and frozen (**Figure 1)**. Since brain cells in the adult brain are highly interconnected and have extensive branches and complex morphology, fresh isolation of cells presents a challenge and often leads to breakage and loss of the distant cellular parts. This may explain the relatively low number of cells in all cell-type specific data sets that were available for LC-MS/MS analysis after FACS sorting **(Supplementary Table 1)**. For LC-MS/MS cell samples were prepared according to mPOP protocol (14). All five types of isolated cell samples from a single control animal resulted in low cell counts, so all samples from this mouse were removed from the follow-up data analysis.

**Figure 1.**
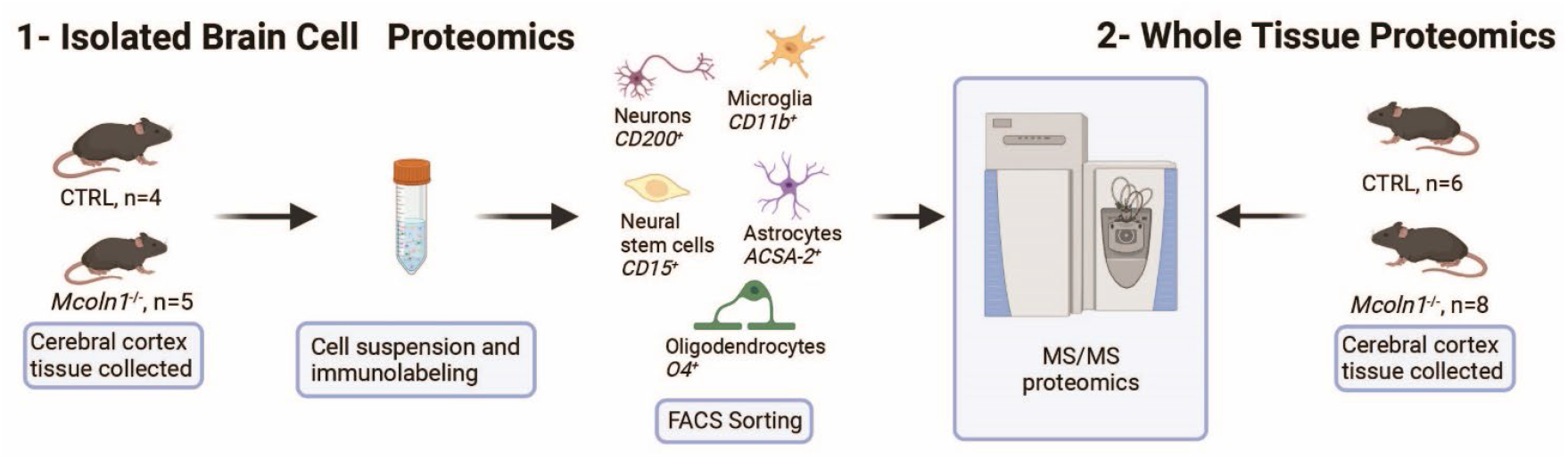
Schematic presentation of proteomics analysis of whole tissue cortical samples and freshly isolated cortical cells in the mouse model of MLIV.

To investigate proteomes in the whole brain tissue, identical parts of cerebral cortex were extracted from the age- and sex-matched *Mcoln1*^*-/-*^ and control mice and snap-frozen on dry ice. Frozen brain tissue was mechanically homogenized, centrifuged and clarified tissue homogenates were digested and eluted peptides were labeled by TMTpro and subsequently separated to 20 fractions. All fractions were dried and, after solubilization in Formic acid, run on mass spectrometer for LC-MS/MS analysis.

### Whole tissue and cell type specific proteomes

LC-MS/MS proteomics analysis identified 7397 proteins from Uniprot mouse and in-house made contaminants databases in the whole tissue cortical homogenates, 446 in neurons, 217 in neural stem cells, 378 in oligodendrocytes, 1985 in astrocytes and 387 in microglia **(Supplementary Table 2)**. All detected non-mouse protein contaminants, as well as mouse albumin and keratin entries were removed from these data sets. Principal component analysis (PCA) separated *Mcoln1*^*-/-*^ samples from controls in all sample sets (**Supplementary Figure 1**). Comparison analysis of whole tissue data set revealed 29 downregulated and 26 upregulated proteins in *Mcoln1*^*-/-*^ cortical tissue as compared to control littermates with log2 foldchange cutoff at 0.6 (foldchange of +/-1.5) and adjusted log10 p<1 (p-value <0.1) **(Figure 2 A and B)**. Among these downregulated proteins, 17 were either myelin or oligodendrocyte lineage enriched proteins (blue asterisks, **Figure 2B**). Remarkably, the whole tissue upregulated proteome in *Mcoln1*^*-/-*^ mice was dominated by lysosomal proteins (16 out of 26) (red asterisks, **Figure 2B**).

**Figure 2.**
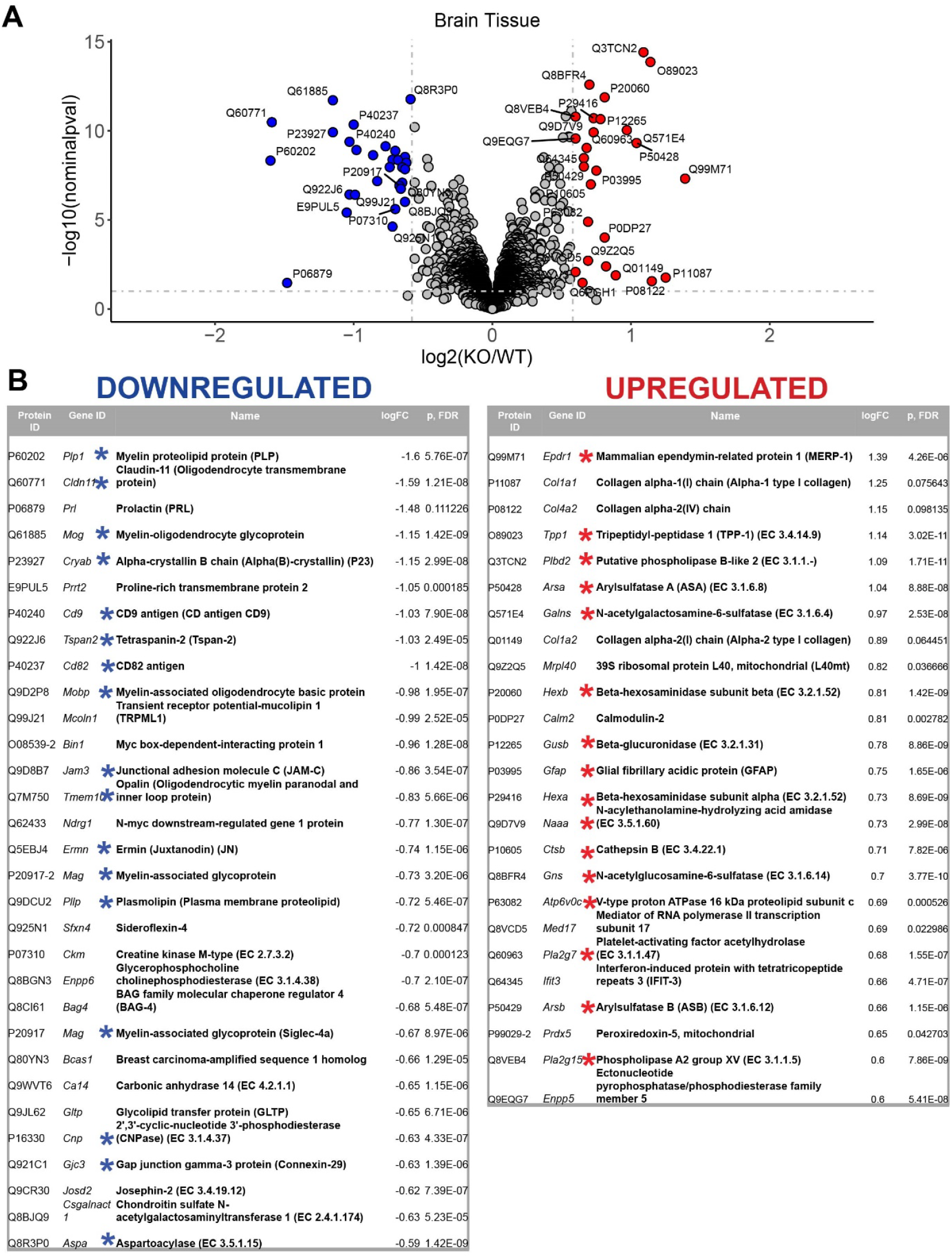
LC-MS/MS proteomics identified upregulated and downregulated proteins in whole tissue cerebral cortex homogenates from *Mcoln1*^*-/-*^ mice. A. Volcano plot showing upregulated and downregulated proteins in whole tissue homogenates from *Mcoln1*^*-/-*^ and control mice; log10 p<0.1, log2FC>0.6. B. List of UP and DOWN regulated proteins in whole tissue of cerebral cortex.

To reveal how loss of TRPML1 affects different brain cell types, we performed comparative analysis of protein abundancies between control and *Mcoln1*^*-/-*^ samples for each cell type **(Figure 3, Supplementary Table 2)**. The highest number of differentially expressed proteins was detected in astrocytes (74 upregulated; 82 downregulated) and microglia (45 upregulated; 47 downregulated), whereas neurons, neuronal stem cells, and oligodendrocytes showed lower number of differential proteins (13, 27, and 13 upregulated; 17, 26, and 5 downregulated per cell type, respectively). We have noticed presence of histones among top five upregulated proteins in neural stem cells and glia, i.e oligodendrocytes, microglia and astrocytes, suggestive of active chromatin reorganization, DNA replication and potentially increased division of the corresponding cell types **(Figure 3)**. Interestingly, despite broad downregulated myelin signature in the whole brain tissue, only a few differentially expressed proteins were detected in the isolated *Mcoln1*^*-/-*^ vs. control oligodendrocytes. Among 17 detected myelin and oligodendrocytes enriched proteins downregulated in the MLIV whole tissue samples, 7 (PLP, MOG, Alpha-Crystallin, MOBP, JAMC, Ermin and CNPase) were resolved in isolated oligodendrocytes but showed no significant changes **(Supplementary Table 2)**. This may indicate that reduced levels of these proteins in whole tissue samples indicate reduced numbers of differentiated oligodendrocytes in the MLIV brain, while isolated O4^+^ *Mcoln1*^*-/-*^ oligodendrocytes didn’t exhibit major cell-intrinsic proteome changes in response to TRPML1 loss.

**Figure 3.**
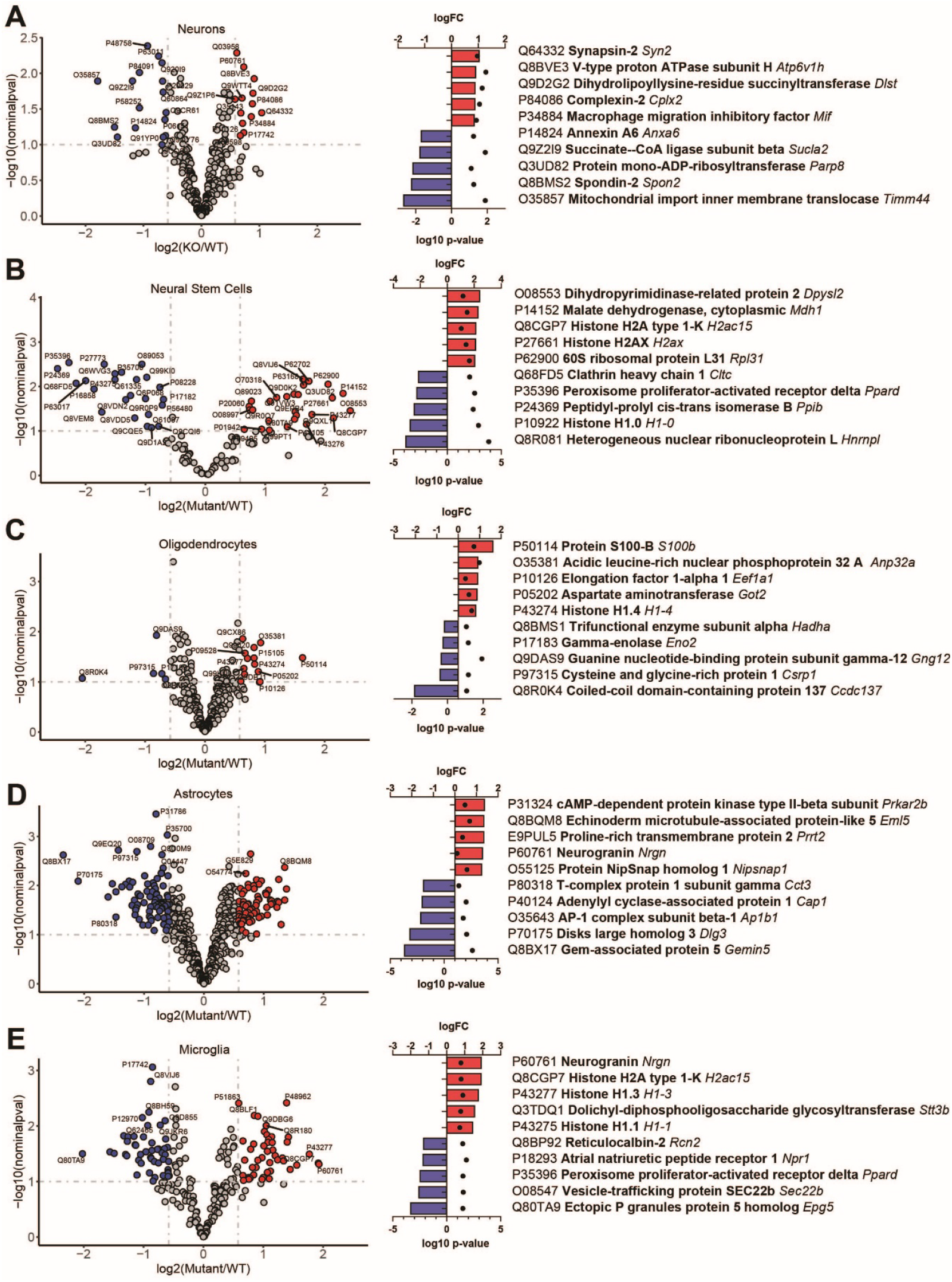
LC-MS/MS proteomics identified upregulated and downregulated proteins in isolated brain cells from *Mcoln1*^*-/-*^ mice. Volcano plot of protein changes in CD200^+^ neurons (A), CD15^+^ neural stem cells (B), O4+ oligodendrocytes (C), ACSA2+ astrocytes (D) and CD11b+ microglia (E) from *Mcoln1*^*-/-*^ cerebral cortex; log10 p<0.1, log2FC>0.6. Left panel show corresponding top 5 upregulated and top 5 downregulated proteins in each data set.

Comparative analysis of overlapping up- and dowregulated proteins between reconstituted whole tissue vs. isolated brain cells proteomes showed no common downregulated and only two common upregulated proteins **(Figure 4A and B)**. Remarkably, the two commonly upregulated proteins are lysosomal enzymes, tripeptidyl-peptidase 1 and beta-hexosaminidase. Upregulation of these lysosomal enzymes across all brain samples confirms that lysosomal dysregulation is a universal cell pathologic mechanism of MLIV.

**Figure 4.**
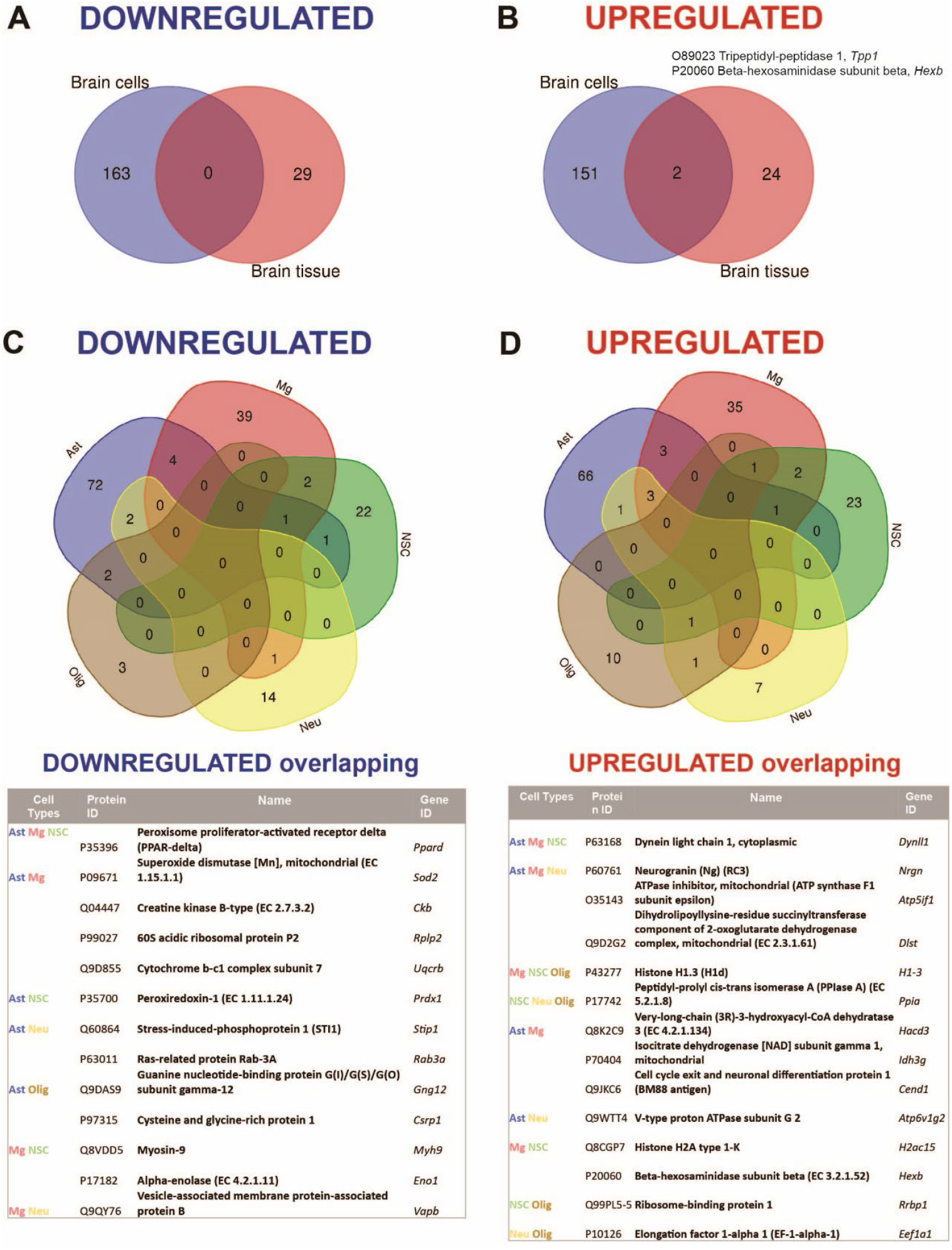
Summary of comparative proteomic analysis showing common and unique findings in whole tissue and isolated brain cells from *Mcoln1*^*-/-*^ mice. A. Venn diagram of common downregulated proteins found in all separate brain cells vs. cortical brain tissue homogenates in *Mcoln1*^*-/-*^ mice. B. Venn diagram of common upregulated proteins found in all separate brain cells vs. cortical brain tissue homogenates in *Mcoln1*^*-/-*^ mice. C. Venn diagram and list of common downregulated proteins found in all separate brain cells isolated from *Mcoln1*^*-/-*^ cerebral cortex. D. Venn diagram and list of common upregulated proteins found in all separate brain cells isolated from *Mcoln1*^*-/-*^ cerebral cortex.

Interestingly, we found very few individual proteins overlapping among isolated neurons, neuronal stem cells, oligodendrocytes, microglia, and astrocytes in either upregulated or downregulated data sets **(Figure 4C and D)**, with the highest number of common up- and downregulated proteins of only 3 and 4, correspondingly, found between microglia and astrocytes. Information on nonoverlapping up- and downregulated proteins for each cell type is available in **Supplementary Table 3**.

### Pathway enrichment analysis

To reconstitute affected biological processes and pathways affected in *Mcoln1*^*-/-*^ brain, we performed gene set enrichment analysis using the Database for Annotation, Visualization and Integrated Discovery (DAVID) on-line portal (https://david.ncifcrf.gov/home.jsp) (15). In line with observations in the comparative analysis of the individual dysregulated proteins, the most significantly enriched downregulated pathways in *Mcoln1*^*-/-*^ cortex in the whole tissue samples were the ones related to white matter development, including myelination, oligodendrocyte differentiation, and gliogenesis **(Table 1)**. The whole list of pathways is available in **Supplementary Table 4**. Tetraspanin CD82 (UniProt P40237) is known to regulate oligodendrocyte migration and differentiation (16, 17) and finding of the InterPro entry IPR018503 Tetraspanin among significantly downregulated terms is in line with general reduction of oligodendrocyte and myelination-related pathways and provides additional insight in mechanisms of the aberrant white matter development in MLIV.

**Table 1.**
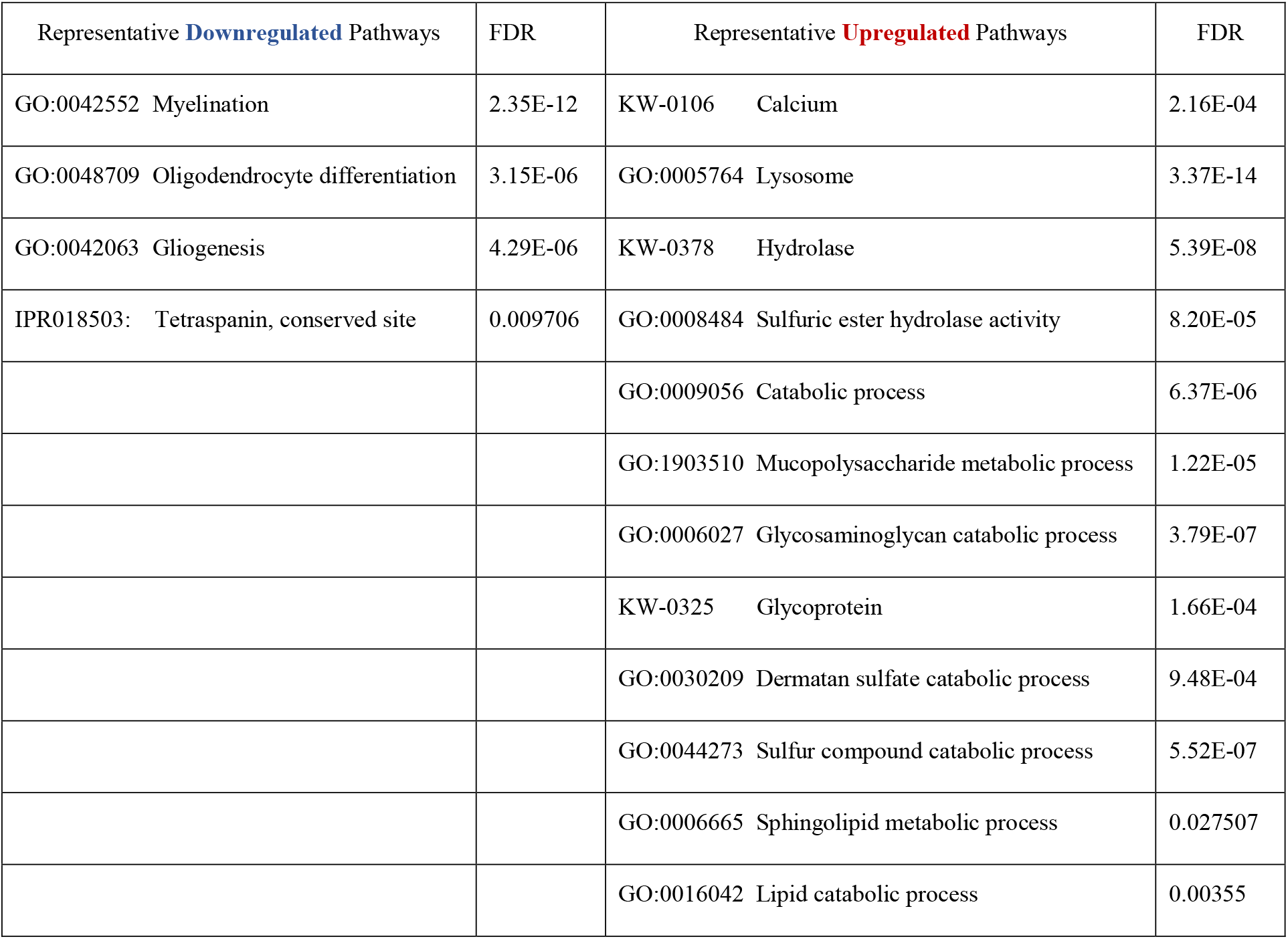
Summary of representative up- and downregulated pathways in whole cortical brain homogenates from *Mcoln1*^*-/-*^ mice.

Upregulated pathways in whole tissue samples included calcium, lysosome/hydrolase and various metabolic and catabolic pathways **(Table 1 and Supplementary Table 4)**. These findings support general knowledge about TRPML1 function as lysosomal calcium channel, confirming on the molecular level that loss of TRPML1 function leads to dysregulated Ca2^+^ signaling and aberrant lysosomal function in MLIV. Enlargement of the lysosomal compartment caused by impaired metabolism and the accumulation of polysaccharides and lipids in lysosomes is a core characteristic of MLIV disease that is reflected in its name, and our data accurately reflect these pathological hallmarks in whole tissue proteome via upregulation of the corresponding cell pathways. At the same time, sulfur-related pathways to our knowledge have not been observed and reported in MLIV brain and provide new insights for future brain pathology evaluation in the brain tissue.

Comparative analysis of the enriched pathways in whole brain vs. brain cells showed that Myelin sheath is the only universal downregulated pathway **(Figure 5A and Supplementary Table 5)**. Seven commonly upregulated pathways included high level structural and metabolic terms such as Cytoplasmic part, Intracellular membrane-bound organelle, Secretory vesicle, Cellular metabolic process, and Lipid metabolism **(Figure 5B and Supplementary Table 5)**.

**Figure 5.**
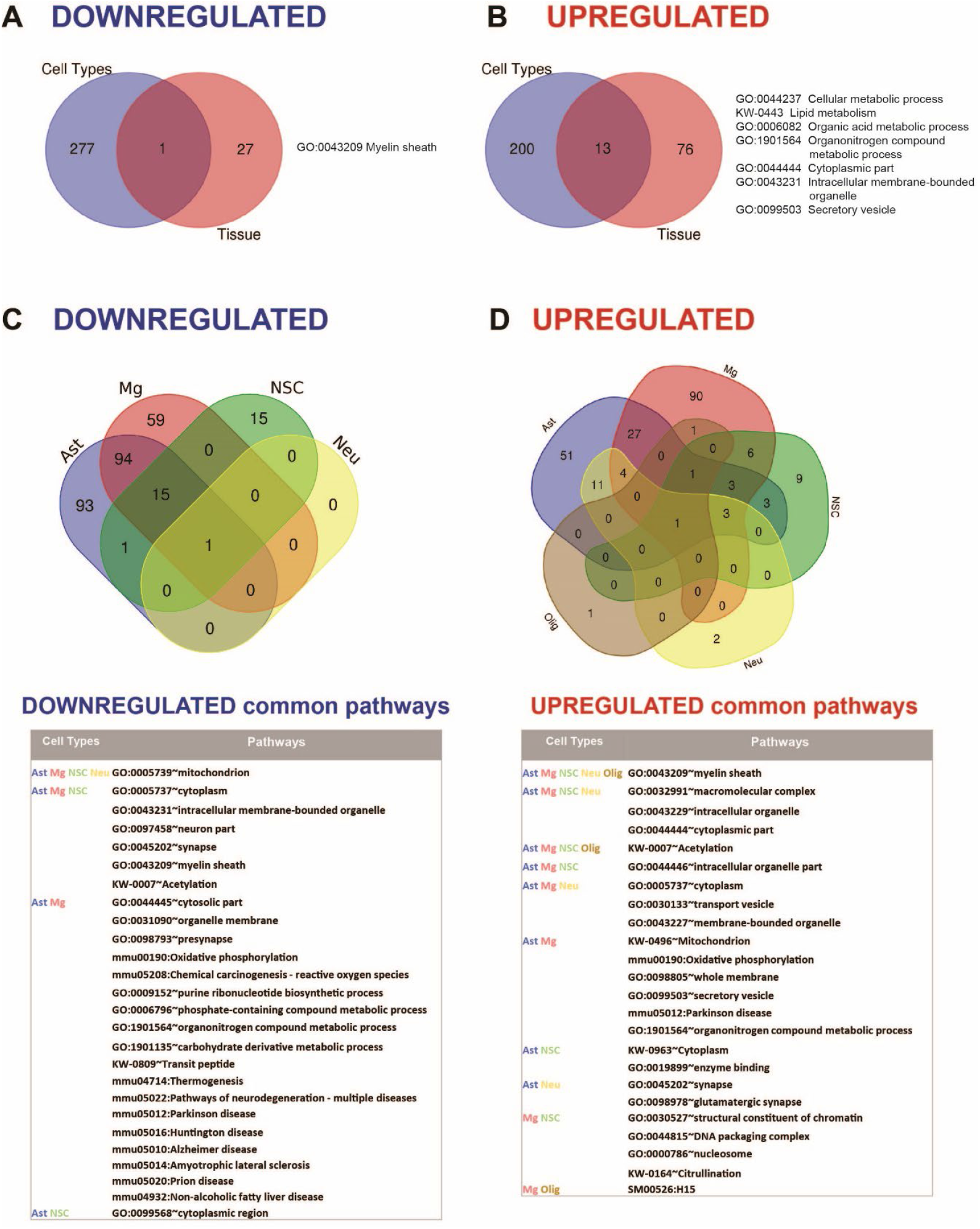
Summary of pathway enrichment analysis showing common and unique findings in whole tissue and isolated brain cells from *Mcoln1*^*-/-*^ mice. A. Venn diagram of common downregulated pathways found in all separate brain cells vs cortical brain tissue homogenates in *Mcoln1*^*-/-*^ mice. B. Venn diagram of common upregulated pathways found in all separate brain cells vs. cortical brain tissue homogenates in *Mcoln1*^*-/-*^ mice. C. Venn diagram and list of representative common downregulated pathways found in all separate brain cells isolated from *Mcoln1*^*-/-*^ cerebral cortex. D. Venn diagram and list of representative common upregulated pathways found in all separate brain cells isolated from *Mcoln1*^*-/-*^ cerebral cortex.

We found no downregulated enriched pathways in isolated oligodendrocytes. Interestingly, Mitochondrion was the only significantly downregulated common term among four other types of isolated brain cells **(Figure 5C)**, and the only downregulated pathway enriched in neurons. The broadest overlap in common downregulated pathways was detected between *Mcoln1*^*-/-*^ astrocytes and microglia, which, together with the high level terms indicating involvement of intracellular compartment and organellar membrane, and previously mentioned mitochondrial energy pathways such as oxidative phosphorylation, also revealed altered various metabolic pathways and pathways of major neurodegenerative diseases such as Parkinson, Alzheimer, Huntington, Prion Diseases, and Amyotrophic Lateral Sclerosis (ALS). This indicates common disease-induced signatures in astrocytic and microglial proteomes in MLIV as well as neurodegenerative settings in general.

Our comparative gene enrichment analysis showed that only one pathway was universally upregulated in all five isolated cell types from *Mcoln1*^*-/-*^ mice, which is Myelin sheath. The total of two upregulated pathways were significantly enriched in either isolated oligodendrocytes or neurons. Besides above-mentioned Myelin sheath, in oligodendrocytes they also included Biosynthesis of amino acids and in neurons Extrinsic component of synaptic vesicle **(Figure 5D and Table 2)**.

**Table 2.**
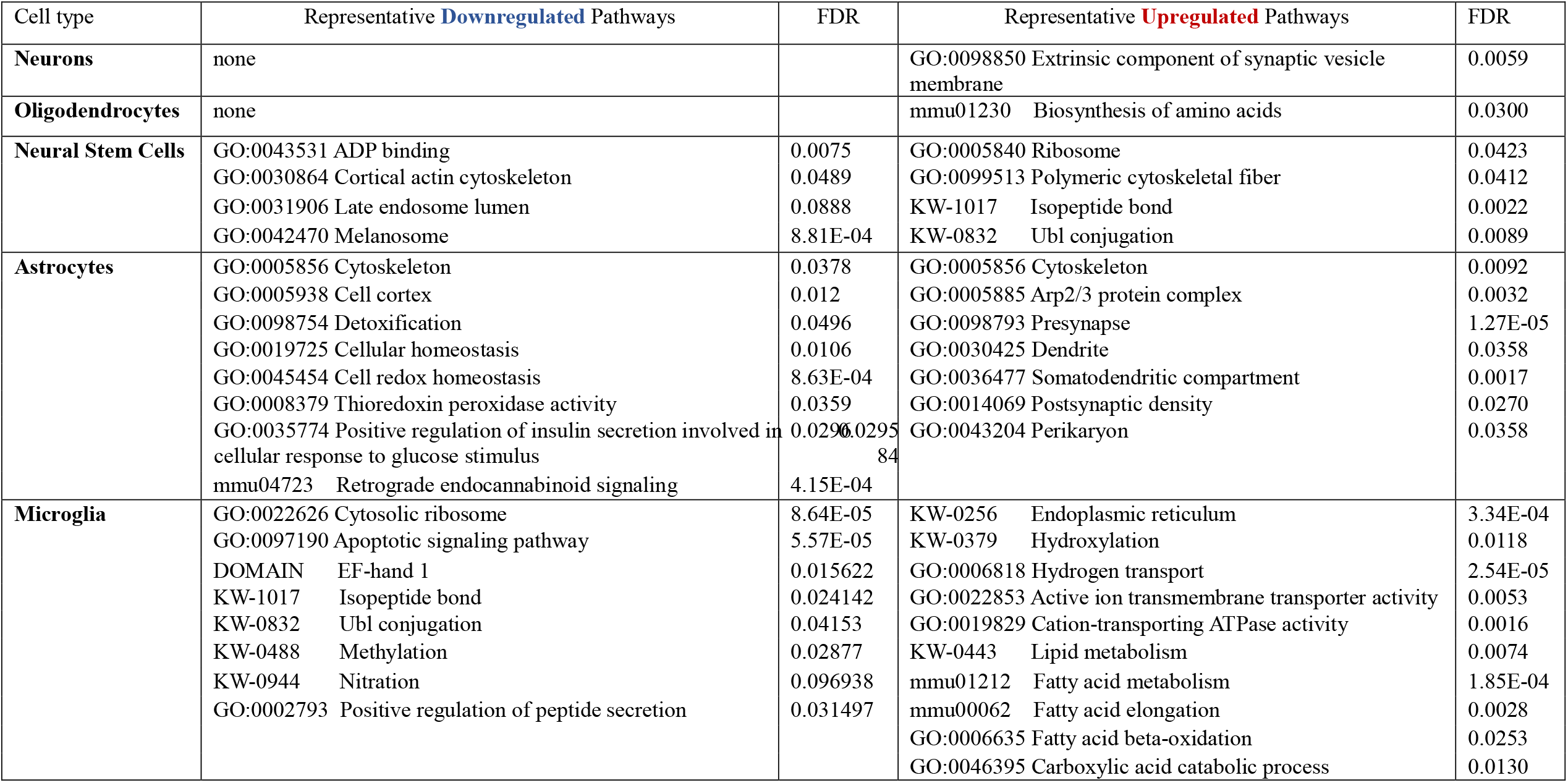
Summary of representative up- and downregulated pathways uniquely identified in isolated brain cells from *Mcoln1*^*-/-*^ cerebral cortex.

Among other interesting findings, we found that Acetylation was upregulated in four isolated cell types, oligodendrocytes being the exclusion. Citrullination was commonly upregulated in microglia and neural stem cells. Other preselected representative overlapping pathways are presented in **(Figure 5D)**, and the full list is available in **Supplementary Table 5**.

On top of common dysregulated pathways in *Mcoln1*^*-/-*^ brain cells, our comparative analysis also revealed unique up- and downregulated pathways in different cell types **(Table 2)**. In astrocytes, Cytoskeleton was dysregulated in both directions, likely indicating reorganization of the astrocyte cytoskeleton while it adopts proinflammatory stimulated, so-called “activated” morphology. In line with this, Cell cortex GO term, describing actin filament and their related proteins network organization beneath plasma membrane in peripheral cytoplasm, was one of the enriched downregulated pathways, whereas actin-related Arp2/3 Protein Complex that facilitates nucleation of branched actin filaments is among enriched upregulated. Other downregulated pathways in astrocytes were related to cell homeostasis, redox stress, and detoxication which, together, manifests the stress and/or “activated” state in *Mcoln1*^*-/-*^ astrocytes. Another cluster of dysregulated pathways in *Mcoln1*^*-/-*^ astrocytes is related to synaptic changes, including Dendrite, Somatodendritic compartment, Post-synaptic as enriched upregulated, and Retrograde endocannabinoid signaling as enriched downregulated pathways.

In microglia, Protein synthesis, Ribosome organization, and Endoplasmic reticulum-related pathways were broadly represented and dysregulated in both directions. Additionally, a cluster of changes in post-translational protein modifications such as Isopeptide bonds, Ubl-conjugation, Nitration, and Methylation was present in downregulated pathways, whereas Hydroxylation was upregulated. Together these findings point towards dysregulated protein synthesis, homeostasis and, likely, degradation in microglia in response to TRPML1 loss. Other unique downregulated pathways in microglia included Ca2^+^-binding EF-hand proteins and Regulation of apoptosis. Lipid metabolism and, more specifically, fatty acid metabolism linked pathways, were broadly present in the unique upregulated group.

In neural stem cells, similar to astrocytes, we have noticed changes in cytoskeleton organization on both up- and downregulated sides. Unique enriched pathways in neural stem cells included Melanosome or Pigment granules, ADP binding, and Late endosome lumen. Ribosome, Protein synthesis, and Post-translational modification pathways were also significantly upregulated; in microglia, these same pathways were downregulated.

Overall, our data provide comprehensive and detailed characterization of the proteome changes in the whole brain tissue and isolated populations of brain cells that reflects previous knowledge of disease pathogenesis and provides new insights into whole brain vs. brain cell proteomes disease mechanisms, by identifying common and revealing unique cell type specific changes in response to loss of the lysosomal channel TRPML1.

## 4 DISCUSSION

Proteomics analysis of the brain has proved to be a powerful tool for enhancing our understanding of its functioning. Transcriptomics approaches have been widely applied across species, stages of development, and diseases to create well-resolved transcriptional brain maps as well as disease-associated transcriptional signatures. However, because transcriptional and proteomic abundances show weak correlation, it is difficult to gain functional insights into brain using only transcriptome data repositories. In recent years, technical advancements in proteomics have helped reconstruct and understand protein composition, expression levels, protein-protein interactions, and post-translational modifications across brain regions and cell types. The purpose of this study was to determine the proteome of the brain in the mouse model of mucolipidosis type IV, at the level of whole tissue as well as in freshly isolated neurons, astrocytes, oligodendrocytes, microglia, and neural stem cells.

Data from our study confirmed known pathophysiological mechanisms of MLIV and revealed previously unknown molecular fingerprints of this disease. Remarkably, cell-type specific proteomics approach allowed to resolve common and cell-type specific proteome changes in *Mcoln1*^*-/-*^ mouse cortex providing deeper insights on disease mechanisms, possible molecular targets, and molecular biomarker candidates for future research and validation. One of the most drastic features of brain pathology in MLIV is reduced myelination and dysmorphic white matter structures, particularly the corpus callosum. This places the disease in the category of hypomyelinating leukodystrophies (18), (19). Like MLIV patients, *Mcoln1*^*-/-*^ mice have a dysgenic corpus callosum (8, 9). Our previous studies showed decreased expression of the mature oligodendrocyte markers myelin-associated glycoprotein precursor (*Mag*), myelin basic protein (*Mbp*), myelin-associated oligodendrocyte basic protein (*Mobp*), and myelin proteolipid protein (*Plp*) in the *Mcoln1*^*-/-*^ cortex during post-natal brain development (20). These myelination markers were stably decreased in *Mcoln1*^*-/-*^ mice in the course of disease (8), indicative of a hypomyelination leukodystrophy rather than a temporary developmental delay in myelination or progressive loss of myelin in the course of disease, known as demyelination. Histopathological analysis in the brain showed that significant thinning of the corpus callosum in MLIV mouse is due to reduced myelination of neuronal fibers and not due to their loss (8, 20), and electron microscopy further confirmed hypomyelination via reduced myelin sheath thickness (9). Reduced expression of the myelination genes *Mag, Mal, Mobp, Mog* and *Cnp* has recently been shown in the cerebral cortex of pre-symptomatic and symptomatic *Mcoln1-/-* mice using RNAseq (21). Importantly, expression of the human myelination genes MOG, MAG, MOBP, LGI4 and CNTNAP1 has also been reported in the human MLIV brain sample. Our whole tissue proteomics analysis in the cortex of *Mcoln1*^*-/-*^ mice clearly demonstrated a similar pattern with the broad range of oligodendrocyte and myelin-enriched proteins showing up as significantly downregulated compared to age- and sex-matched control mice **(Figure 2)**. Gene enrichment analysis showed a big cluster of myelination, oligodendrocyte differentiation, and axon ensheathment pathways significantly enriched and downregulated **(Figure 5 and Supplementary Table 4)** with GO:0043209 Myelin sheath being the only common downregulated pathway among brain tissue and isolated brain cells proteomes, fully supporting our previous findings. Interestingly, although many of the oligodendrocyte signature proteins were resolved in the isolated oligodendrocytes, unlike in whole tissue samples, they did not show reduced abundance. Reduction in myelin and oligodendrocyte protein abundancies in whole tissue indicates fewer oligodendrocytes and less deposited myelin in the *Mcoln1*^*-/-*^ brain. However, despite being less populated in MLIV brain, differentiated and isolated *Mcoln1*^*-/-*^ oligodendrocytes did not show drastic changes in their proteomes as compared to control brain oligodendrocytes, suggesting that deficient differentiation of oligodendrocyte precursors to myelinating oligodendrocytes may be largely responsible for hypomyelination. In summary, while the detailed mechanisms of TRPML1 role in brain myelination is not fully understood and requires further investigation, our data support clinical observations in MLIV patients and previously obtained data in the *Mcoln1*^*-/-*^ mouse model and confirm the hypomyelinating leukodystrophy phenotype.

Another drastic feature of brain pathology in *Mcoln1*^*-/-*^ mice and human MLIV brain is the enlargement of the lysosomal compartment and the accumulation of the lysosomal “storage bodies”, or inclusions, that are structurally and biochemically heterogeneous and contain membranous and granular electron-dense storage due to accumulation of undegraded lipids, polysaccharides, and proteins (9, 22, 23). Our data confirm these observations on proteomics level. We report broad upregulation of lysosomal enzymes and structural lysosomal proteins in whole tissue homogenates from *Mcoln1*^*-/-*^ mice, with two of the detected lysosomal enzymes commonly upregulated in whole brain and isolated brain cell data sets **(Figures 2 and 4B)**. Using various techniques, including transcriptomics, immunohistochemistry, and enzymatic activity assays, upregulation of lysosomal proteins was commonly reported in other lysosomal storage disorders, such as NPC, Gaucher, mucopolysaccharidoses, and other, and is thought to represent a mechanism to compensate for impaired lysosomal function (24, 25). A similar increase of lysosomal proteins was reported in the single MLIV brain autopsy proteome, highlighting the universal nature of these observations in the brain tissue across species (21).

We also observed upregulation of glycoproteins in *Mcoln1*^*-/-*^ cortex (**Table 1**). A closer look revealed that many of these are lysosomal hydrolyzes. Interestingly, we also found a group of glycosylated collagens Col1a1, Col1a2 and Col4a2 significantly elevated in *Mcoln1*^*-/-*^ whole cortex samples. Collagens are a key component of the extracellular matrix (ECM) in the brain and are primarily produced by astrocytes. Some evidence suggests that collagens IV and I may also be produced and play a role in oligodendrocyte development and function, i.e. adhesion, migration and differentiation of oligodendrocytes during development (26). We did not detect presence of collagens in either isolated astrocytes or oligodendrocytes. Elevated levels of collagens have been reported in neurodegenerative diseases, including Parkinson’s (27) and multiple sclerosis (28, 29). Increased vascular collagen IV has been linked to Alzheimer Disease (30) and amyotrophic lateral sclerosis (ALS) (31). Accumulations of collagens have also been reported in response to inhibition of autophagy, or more specifically, endoplasmic reticulum (ER)-phagy, a process in which excess of collagens is being removed via selective targeting for degradation on lysosomes (32, 33) and is regarded as marker of ER-phagy. Hence, an observed increase in collagens may point to inhibition of this form of autophagy in MLIV, a likely additional consequence of lysosomal function disruption due to loss of TRPML1. Overall, the mechanism leading to increase of collagens in MLIV is unknown and demands further scientific elucidation.

In line with previous reports of lysosomal pathology and lysosomal storage composition, together with the higher abundancies of lysosomal proteins, our pathway analysis on whole brain tissue also showed upregulation of the metabolic and catabolic pathways of lipids and polysaccharides.

Importantly, lipid metabolism was one of 7 pathways universally upregulated between brain tissue and all isolated cell types. On top of shared presence of lipid metabolism dysregulation across brain tissue and brain cell samples, proteome analysis of isolated cell types provided deeper insights, such as upregulation of fatty acid metabolism-related pathways unique to microglia.

Another group of cellular pathways consistently dysregulated in all isolated brain cells, except oligodendrocytes, is related to mitochondrion function and oxidative phosphorylation **(Figure 5)**. Mitochondrion-related dysregulated pathways were very broadly presented in our analysis, particularly in microglia and astrocytes, and, together with the above-mentioned general mitochondrion and oxidative phosphorylation terms, included some more specific structural terms such as mitochondrial envelope, inner membrane, outer membrane, or more narrow parts of the redox chain, such as ATPase or NADH dehydrogenase complex **(Supplementary Table 4)**. Interestingly, while mitochondrial aberrations in the form of fragmentation and decreased mitochondrial Ca^2+^ buffering capacity were reported in MLIV patient fibroblasts (34), analysis of electron micrographs in Shaeffer collaterals of hippocampus did not report similar structural changes in the *Mcoln1*^*-/-*^ mouse brain (9). To our knowledge, this is the first report of mitochondrial changes suggestive of aberrant mitochondrial function in MLIV mouse. In the literature, presence of fragmented mitochondria in MLIV fibroblasts was hypothesized to be caused by insufficient mitophagy, a process of selective degradation of senescent mitochondria on lysosome as a result of impaired lysosomal function due to loss of TRPML1 function (34). Later, a mechanism of lysosomal-mitochondrial communication was established, where TRPML1 was shown to facilitate direct transfer of Ca^2+^ from lysosomes to mitochondria (35). Loss of TRPML1 function in MLIV fibroblasts lead to lower mitochondrial Ca^2+^, reduced lysosome-to-mitochondria contact-dependent transfer of Ca^2+^, and altered lysosome-mitochondrial contact tethering dynamics, with more lysosomes involved and longer tethering contacts in MLIV cells. Authors suggested that lysosome-to-mitochondria TRPML1-mediated Ca^2+^ influx can be important to facilitate downstream calcium-dependent mitochondrial functions, including oxidative phosphorylation, motility, and reactive oxygen species (ROS) signaling. Therefore, there is a possibility that either impaired turnover of mitochondria or defective lysosome-mitochondria Ca^2+^ transfer caused by loss of TRPML1 function in MLIV may contribute to the broad mitochondrial dysregulation we observed in whole brain tissue and cell-type specific data sets in this study.

Despite the broad overlap in mitochondrial changes between cell types, mitochondria-related pathways were not enriched in whole brain tissue samples. This may be due to low mitochondrial protein sensitivity in whole tissue samples and saturation of the MS detector by proteins that are abundant in brain tissue. This example illustrates the advantage of cell type specific versus whole brain proteomics approach to resolve pathophysiologic changes.

Proinflammatory activation of microglia and astrocytes, often referred to as microgliosis and astrocytosis, is a persistent feature of brain pathology in MLIV that is evident in MLIV mouse and human brain tissue starting from the early postnatal (mouse MLIV) and late prenatal (human MLIV) stages of development to end-stage of disease (4, 9, 22, 36, 37). Increased levels of proinflammatory cytokines and chemokines previously reported in MLIV cortical homogenates and astrocyte cultures provide evidence of neuroinflammatory axis in MLIV (37). Despite this, we previously found that AAV9-mediated gene transfer of *MCOLN1*/TRPML1 directed to neurons was sufficient to fully restore neurologic function in *Mcoln1*^*-/-*^ mice without reducing microgliosis and astrocytosis, therefore dismissing the primary role of these cells in driving MLIV pathogenesis (38). Studies of *Mcoln1*^*-/-*^ microglia showed decreased expression of microglial surface markers CX3CR1 and CD11b and elevation of markers associated with activation of microglia such as CD86 and MHCII, increased free radical content, expression of iNOS, and shift to glycolytic metabolism (36). Transcriptional profiling of freshly isolated CX3CR1^+^CD11b^+^CD45^+^ microglia isolated from MLIV mice with subsequent pathway analysis demonstrated changes in lysosomal function, immune cell activation, and neurodegeneration-related gene sets (36). Notably, we also see enrichment of neurodegenerative disease-related pathways in microglia in our study. CD86, HIF1a and HIF1a target genes were not resolved in our microglia samples sets. We also did not find immune signaling signature proteins in microglial data sets in our study, including CCL5, which was reported as one of the most significantly overexpressed gene in (36). These differences can be explained by multiple factors including fewer than optimal to see low abundance proteins starting peptide load due to low cell numbers in original samples, the use of different protocol and antigen markers for microglia isolation resulting in different population of microglia subsets undergoing analysis, lower protein coverage in our study than transcript coverage in (36), as well as limited translatability between transcriptomic and proteomic abundancies. Our data, however, revealed that, together with commonly dysregulated pathway signature including membrane organization, secreted vesicles, mitochondria, lipid metabolism, etc., some pathways clusters, such as apoptosis signaling and cell death, dysregulation of protein synthesis, multiple post-translation modifications alternations, and upregulation of fatty-acid metabolism related pathways, were a distinct feature of *Mcoln1-/-* microglia.

This study provides a protein fingerprint of MLIV brain disease in *Mcoln1*^*-/-*^ mice, confirming previously known pathological hallmarks and adding new depth to the understanding of the molecular changes and pathways underlying the disease. Both LC MS/MS proteomics approaches, using either whole tissue or isolated cell type-specific fractions, were synergistic in providing in-depth information on the proteome signature of MLIV. The major observations at the level of whole brain tissue were downregulation of myelination-related pathway, and upregulation of calcium, lysosome, and metabolic pathways, including lipids and carbohydrates. While analyzing proteomes of *Mcoln1*^*-/-*^ neurons, neural stem cells, astrocytes, microglia, and oligodendrocytes, we found commonly dysregulated pathways between these cell types that included organellar membrane, membrane protein complexes, mitochondria and oxidative phosphorylation, synapse composition, lipid metabolism, chromatin organization, secretory or transport vesicles, and some post-translational protein modifications. This portfolio of pathways creates a signature of the basic, or common, alternations in cells in response to loss of TRPML1 function. Additionally, our analysis also showed cell-type specific responses to loss of TRPML1 that might be especially insightful for understanding the role of this lysosomal channel in cell type specific setting. We found that among all studied cell types, despite strong hypomyelination phenotype in MLIV, isolated O4+ *Mcoln1*^*-/-*^ oligodendrocytes had minimal changes at the level of proteome when compared to oligodendrocytes from the age- and sex-matched controls. Major changes in neurons were found in proteins related to mitochondria, myelin sheath, and synaptic membrane organization. In addition to this, in neural stem cells, pathways related to melanosomes and protein synthesis were enriched. Yet, the highest number of dysregulated pathways were found in microglia and astrocytes, that, together with neurodegeneration-related, membrane organization, metabolic and mitochondria-related pathways commonly enriched in both populations, also showed a signature of distinct, specific changes. In astrocytes, these changes included cytoskeleton-, cell homeostasis-, endocannabinoid signaling-, and postsynaptic membrane-related pathways, whereas microglia showed enriched protein synthesis, endoplasmic-reticulum-related, fatty acid metabolism, and regulation apoptosis pathways.

Together with resolved whole tissue and cell type-specific proteome profile of MLIV, our data provide a helpful insight into potential molecular biomarkers of MLIV. Since TRPML1 is a transmembrane lysosomal channel, in-vitro assay reporting restoration of its function for future efficacy and potency assay development has to rely on direct measurement of its ion conduction which presents technical challenges. Even more importantly, easily accessible surrogate biomarkers that could report activity of TRPML1 in the brain in accessible tissues, such as serum or plasma, for translational and interventional studies are currently missing. Our study provides insight into potential molecular biomarkers for future validation. For example, based on our data, hexosaminidase B (HexB) and tripeptidyl-peptidase 1 (TPP1) could be possible biomarker candidates. Both proteins are lysosomal enzymes with increased abundances in response to TRPML1 loss in both brain tissue and brain cells data sets. This suggests either their compensatory upregulation in response to lysosomal dysfunction or reflects enlargement of the lysosomal compartment due to impaired lysosomal turnover. Future studies will be required to validate the use of HexB, TPP1 or other lysosomal proteins as biomarkers for MLIV.

## 8 Author Contributions

BB and YG contributed to conception and design of the study, data interpretation and wrote manuscript. BB, MS, MCC, ML and ZN performed experiments, SS performed data analysis for this study, MS and SS contributed to manuscript writing. BB created a data repository. All authors contributed to manuscript revision, read, and approved the submitted version.

## 10 Acknowledgments

The authors are thankful to Dr. Albert Misko for thought-provoking discussions and big contribution to understanding natural history of MLIV and to the Mucolipidosis Type IV (ML4) Foundation and its executive director, Dr. Rebecca Oberman, for support of research and tireless efforts to accelerate therapy development and translationally oriented research for MLIV.

## 12 Data Availability Statement

The datasets GENERATED and ANALYZED for this study can be found in the MassIVE data repository https://massive.ucsd.edu/ProteoSAFe/dataset.jsp?task=1ecdab485bf046b5b8457e71750b5eac All raw data can be downloaded here: ftp://MSV000091824@massive.ucsd.edu

